# Oxicam-type NSAIDs enhance *Agrobacterium*-mediated transformation in plants

**DOI:** 10.1101/2020.12.15.422982

**Authors:** Seung-won Choi, Kie Kumaishi, Reiko Motohashi, Harumi Enoki, Wiluk Chacuttayapong, Tadashi Takamizo, Hiroaki Saika, Masaki Endo, Tetsuya Yamada, Aya Hirose, Nobuya Koizuka, Seisuke Kimura, Yaichi Kawakatsu, Hiroyuki Koga, Emi Ito, Ken Shirasu, Yasunori Ichihashi

## Abstract

*Agrobacterium*-mediated transformation represents a key innovation for plant breeding and is routinely used in research and applied biology. However, for several species, the efficacy of transformation is limited. In this study, we discovered that oxicam-type nonsteroidal anti-inflammatory drugs (NSAIDs), including tenoxicam (TNX), enhance the efficiency of *Agrobacterium*-mediated transient transformation in the model species *Arabidopsis thaliana* via leaf infiltration and can be successfully applied in analyses of the subcellular localisation of fluorescent fusion proteins. TNX acts as an inhibitor of plant immune responses and lacks similar transient transformation efficiency in a *dde2/ein2/pad4/sid2* quadruple mutant background, thereby indicating that TNX increases the efficiency of *Agrobacterium* infection via a transient shutdown of the immune system mediated by jasmonic acid, ethylene, and salicylic acid signalling. In addition, we found that TNX enhances the efficiency of stable transformation in crops of agricultural and economic importance, such as Jatropha and maize, indicating that TNX can enhance the integration of exogenous DNA into the plant genome via an increased introduction of DNA into plant cells. Given that treatment with oxicam compounds is simple, cost effective, and has broad utility, we anticipate that this discovery will contribute to accelerating genome-editing technologies in plants.

## Introduction

The continual increase in global population poses considerable challenges with respect to maintaining the overall sustainability of food production, and in this regard, the genetic modification of crops via plant transformation is seen as providing one solution to cope with prospect of future food insecurity. Owing to advances in sequencing technology, whole-genome sequences of many plant species, including crops, have been determined, thereby enabling comprehensive genetic analyses (The Arabidopsis Genome Initiative, 2000; Goff et al., 2002; Yu et al., 2002). In addition, recent technical advances in genome editing, such as TAL effector nucleases and CRISPR/Cas, are facilitating the generation of targeted DNA double-strand breaks with high specificity in complex genomes. Therefore, technically, the manipulation of plant genomes via genetic transformation is becoming increasingly viable. However, in the case of many plant species, including crops of agricultural and economic importance, current modification approaches remain inefficient. Accordingly, in order to maximise the potential implementation of technological knowledge to feed the world, it is essential to further enhance the capacity and efficiency of plant transformation (Altpeter et al., 2016).

Common soil bacteria in the genus *Agrobacterium* have been found to have the unique ability of interkingdom genetic transfer, notably, the incorporation of transfer DNA (T-DNA) segments into plant genomes (Escobar and Dandekar, 2003; Tzfira et al., 2004). This ability of *Agrobacterium* species to transfer DNA to plants has been extensively exploited for genetic engineering purposes by replacing oncogenes within bacterial T-DNA with any gene of interest. *Agrobacterium*-mediated genetic transformation has become a routine procedure in basic plant research as well as a principal means of generating transgenic plants for the agricultural biotechnology industry (Gelvin, 2005). However, some plant species show heightened immune responses to suppress *Agrobacterium*-mediated transformation. For example, it is well established that ethylene (ET)- and salicylic acid (SA)-mediated immune responses are deployed by plants to restrict *Agrobacterium*-mediated transformation (Gaspar et al., 2004; Yuan et al., 2007; Anand et al., 2008; Nonaka et al., 2008b, 2008a; Lee et al., 2009). Indeed, immunity-related mutants and transformants have been reported to enhance the efficacy of *Agrobacterium*-mediated transient transformation [e.g. *dde2/ein2/pad4/sid2* (Tsuda et al., 2009) and *NahG* (Rosas-Díaz et al., 2017)]. As an alternative strategy, transformation efficiency can also be enhanced by modifying the virulence of *Agrobacterium*, which influences plant immunity (Hiei et al., 1994; Ishida et al., 1996; Piers et al., 1996; Rashid et al., 1996; de Groot et al., 1998; Wenck et al., 1999; Rohini and Rao, 2000; Kunik et al., 2001; Komari et al., 2006; Kimura et al., 2015). Accordingly, controlling the immunity responses of target plants is considered a key strategy that could be used to address current inefficiencies in plant transformation.

During the course of previous chemical screening (Noutoshi et al., 2012), we established that oxicam-type nonsteroidal anti-inflammatory drugs (NSAIDs), such as tenoxicam (TNX), can function as plant immune inhibitors (Ishihama et al., 2020). In the present study, we report that brief treatment with immune-inhibiting oxicams is effective in enhancing transformation efficiency in several plant species, including agricultural crops. Although the effect of these chemicals appears to be tissue specific, we believe that this simple approach, entailing the use a transient chemical treatment, has considerable potential to facilitate the practical application of technological principles for the purposes of plant transformation.

## Results and Discussion

### Oxicam treatment enhances the efficiency of *Agrobacterium*-mediated transient transformation of *Arabidopsis* leaves

The suppression of plant immune responses represents a key strategy for enhancing the efficiency of *Agrobacterium*-mediated transient transformation. In *A. thaliana*, agroinfiltration of leaves has been used to characterise gene functions in several mutants and transformants defective in immune responses (Tsuda et al., 2009; Sardesai et al., 2013; Rosas-Díaz et al., 2017). However, this approach would not be a suitable system for investigating the functions and subcellular localisation of unknown genes, as it does not exclude the possibility that such mutations may affect the functions of these genes. On the basis of the findings of our previous study, indicating that tenoxicam (TNX) inhibits plant immunity [Fig. 1; Ishihama et al. (2020)], we sought in the present study to determine whether a brief treatment of an *Agrobacterium* suspension with TNX would enable us to utilise wild-type *A. thaliana* for agroinfiltration in a manner similar to immunity-deficient mutants. To assess the feasibility of this approach, we added TNX to a suspension culture of *35S::GUS*-carrying *Agrobacterium* prior to infiltration as a brief treatment, and compared the efficiency of transient transformation when the same culture without TNX treatment was used to agroinfiltrate the quadruple mutant (*dde2/ein2/pad4/sid2*), defective in ET-, SA-, and jasmonic acid (JA)-mediated immunity, which displays high efficiency (Tsuda et al., 2009). We accordingly observed that TNX-treated wild-type plants showed markedly enhanced GUS activity, which was seven times higher than that in plants transformed with non-treated *Agrobacterium*, and established that this heightened activity was equivalent to that observed when using non-treated *Agrobacterium* to infiltrate the mutant leaves (Fig. 2A and B). These findings thus provide evidence that the efficiency of DNA transformation was elevated owing to an increased efficiency in infection achieved by reducing plant immunity via a brief treatment with TNX. We subsequently examined the subcellular localisation of expressed marker proteins, including nuclear-localised histone 2B (H2B-GFP), the *trans*-Golgi network (TGN) marker Venus-SYP61, and plasma membrane-bound aquaporin (PIP2a-mCherry) following infiltration with TNX-treated *Agrobacterium*. We accordingly observed the anticipated localisation of these proteins in both TNX-treated and non-treated leaves (Fig. 2B), indicating that transient application of TNX can be effectively used to monitor the subcellular localisation of proteins. In contrast, however, TNX treatment appeared to have no similar effects with regards to root and flower transformation (Table S1 and S2). To determine the immune-related pathways targeted by TNX in recipient plants, we subjected the *dde2/ein2/pad4/sid2* quadruple mutant to transfection using TNX-treated *Agrobacterium*, and found that the efficiency of transient transformation, as indicated by GUS activity, in the quadruple mutant was similar to that in the mutant infiltrated with non-treated cells (Fig. 2C). These observations thus indicate that TNX can effectively inhibit the signalling pathways mediated by SA, JA, and ET. In addition to TNX, we also examined the effects of five other oxicams, namely, meloxicam (MLX), piroxicam (PRX), ampiroxicam (APRX), sudoxicam (SDX), and lornoxicam (LNX, Fig. 1A), on *Agrobacterium*-mediated transient transformation of *Arabidopsis* leaves, and accordingly observed that all five increased GUS activity. Indeed, LNX and SDX were found to be significantly effective, similar to TNX (Fig. 2D). These results thus indicate that transient oxicam treatment represents a highly effective new approach for enhancing the efficiency of leaf agroinfiltration in wild-type *A. thaliana*.

**Figure 1.**
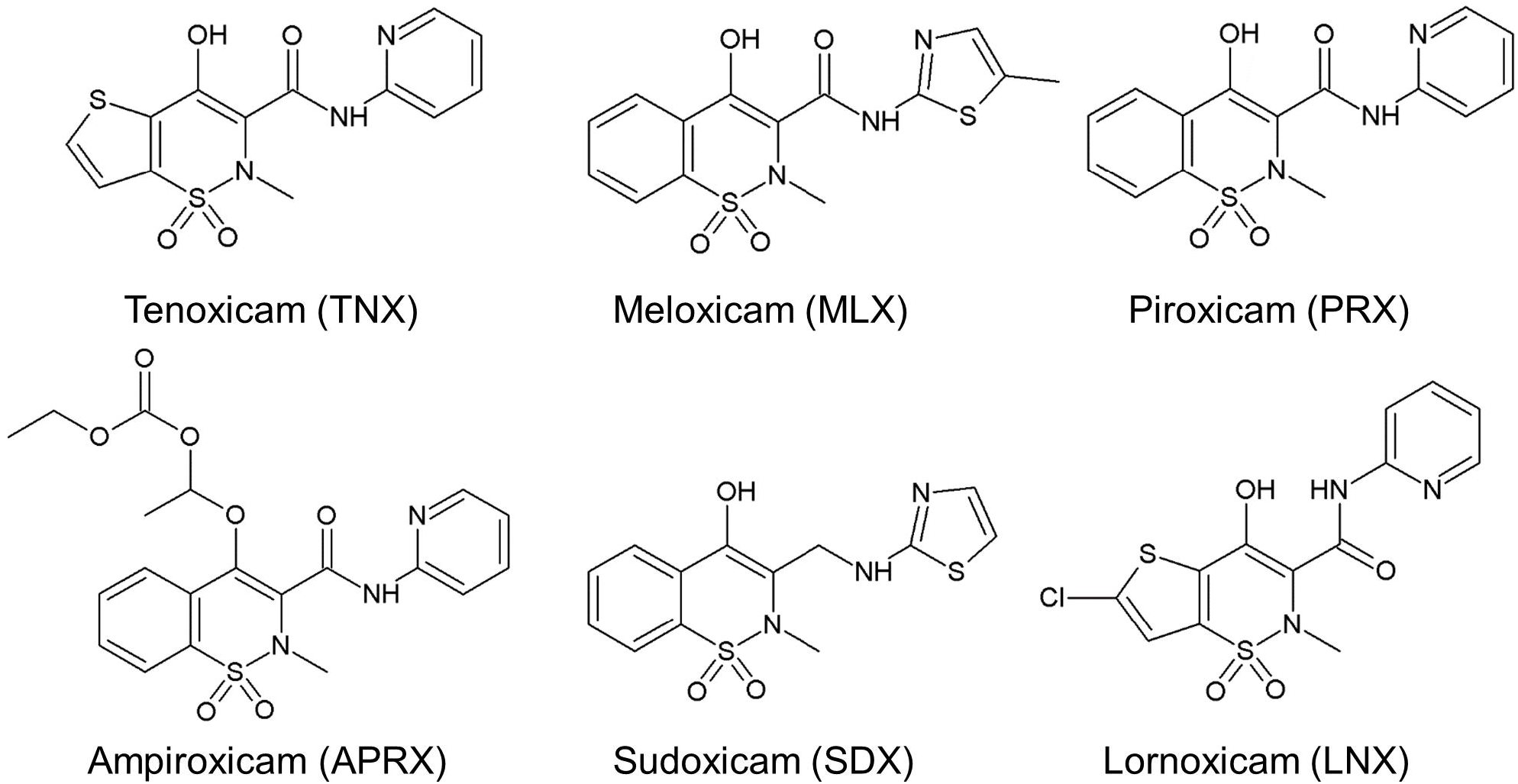
The structures of oxicam compounds. Tenoxicam (TNX), meloxicam (MLX), piroxicam (PRX), ampiroxicam (APRX), sudoxicam (SDX), and lornoxicam (LNX).

**Figure 2.**
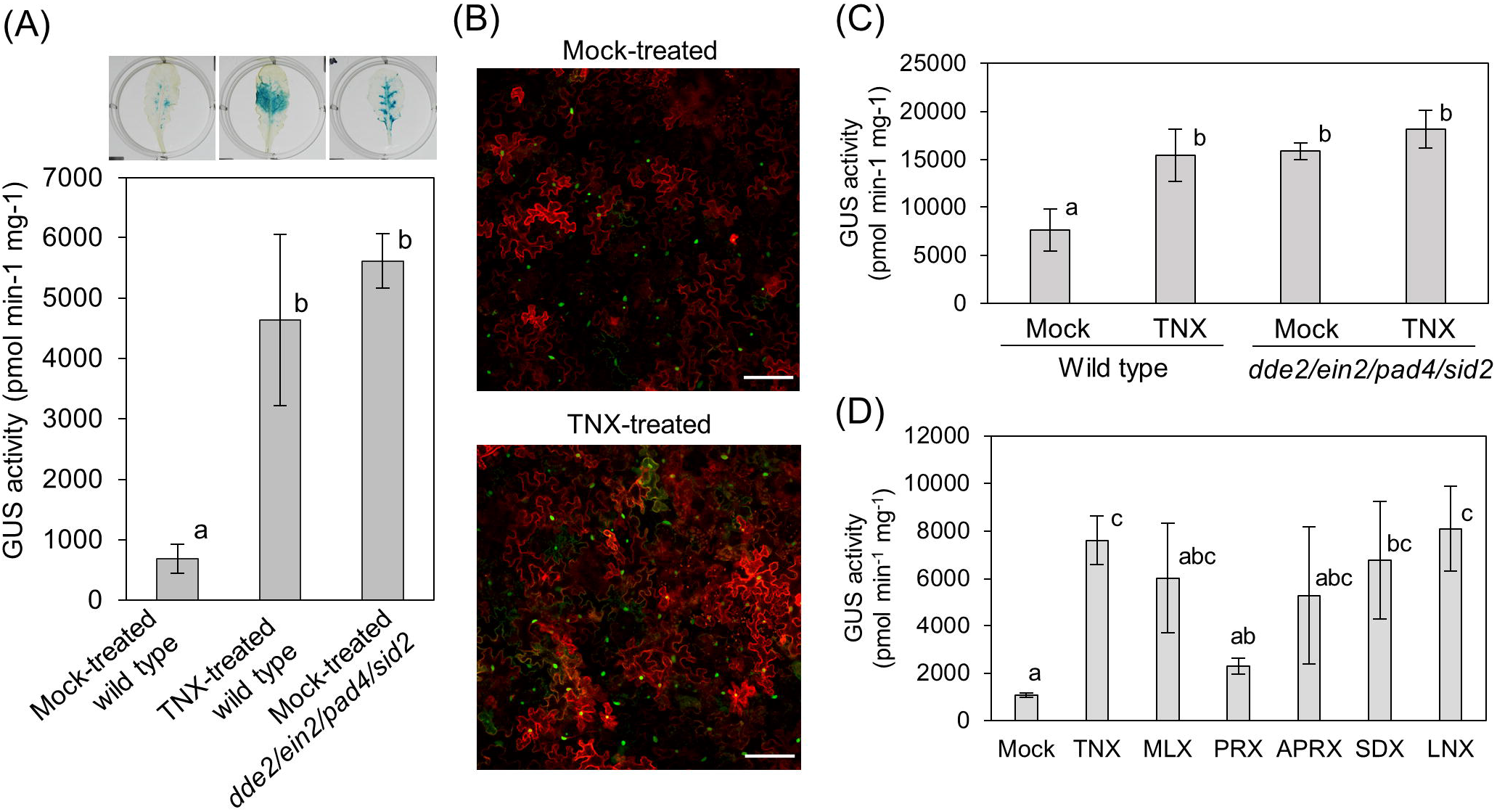
Oxicam treatment enhances the efficiency of *Agrobacterium*-mediated transient transformation in *Arabidopsis thaliana*. (A and B) Leaves of wild-type and *dde2/ein2/pad4/sid2* quadruple mutant *Arabidopsis thaliana* were mock- and tenoxicam [TNX (100 μM)]-treated and inoculated with *Agrobacterium* carrying a *35S::GUS*. (A) After 2 days, GUS enzyme activity was determined by histochemical staining and a quantitative MUG assay. The blue colour indicates GUS activity. Experiments were repeated five times with similar results. Bars represent the means and standard errors of data obtained from three biological replicates. (B) Transgenic leaves inoculated with *Agrobacterium* strains carrying the constructs *RPS::H2B-GFP* for the nucleus, *35S::Venus-SYP61* for the *trans*-Golgi network, and *35S::PIP2A-mCherry* for the plasma membrane in the mock- or TNX (100 μM)-treated wild-type plants. Fluorescence signals in epidermal cells were observed 2 days after infiltration. Bars = 100 μm. (C and D) Quantitative MUG assay to assess the effect of TNX on quadruple mutant plants (C) and the effect of oxicams on wild-type plants (D). Agroinfiltration conditions were the same as those in (A). Different letters indicate statistically significant differences, as determined using Tukey’s test (*P* < 0.05).

### TNX can be applied for the transformation of agricultural crops

On the basis of our finding that transient treatment with TNX can enhance the efficiency of *Agrobacterium*-mediated DNA transformation of the vegetative shoot tissues of *A. thaliana* (Fig. 2), we anticipated that similar treatment might also enable us to enhance the efficiency of transformation of the vegetative tissues of agricultural crops. To verify this possibility, we evaluated the effect of TNX on *Agrobacterium*-mediated transformation of Jatropha (*J. curcas*) and maize, as the transformation methods used for these species typically utilise vegetative tissues such as cotyledon explants and immature embryos, respectively [Fig. 3A-B, Fig. 4A; Ishida et al. (1996); Enoki et al. (2017)]. Given that Jatropha is an oil seed plant that yields biofuel, and that the seed oil can be readily processed as a replacement for petroleum-based diesel fuel (Fig. 2A, Forson et al., 2004), it would be desirable to develop techniques that can be used to enhance the transformation efficiency of Jatropha for industrial applications. In the present study, we performed Jatropha transformation by adding TNX to the co-cultivation suspension, and to determine the stability of transformation, we introduced selection vectors that enabled us to evaluate the transformation rate by PCR genotyping for the transgenes (Fig. 2C). We accordingly found that upon TNX treatment the rate of stable transformation with respect to the number of transformed soil-acclimated plants (11.5%–18.8%) was approximately 4- to 16-fold higher than that of the control plants (1.15%–2.63%, Fig. 3C). Similarly, when we calculated the transformation rate with respect to the number of cotyledon explants as starting materials, the rate was still 8-fold higher in the TNX treatment (0.285%) than that in the control (0.033%) for experiment 1 (Fig. 2C). Collectively, these data indicate that pre-treatment with TNX can significantly enhance the efficiency of stable *Agrobacterium*-mediated transformation of Jatropha.

**Figure 3.**
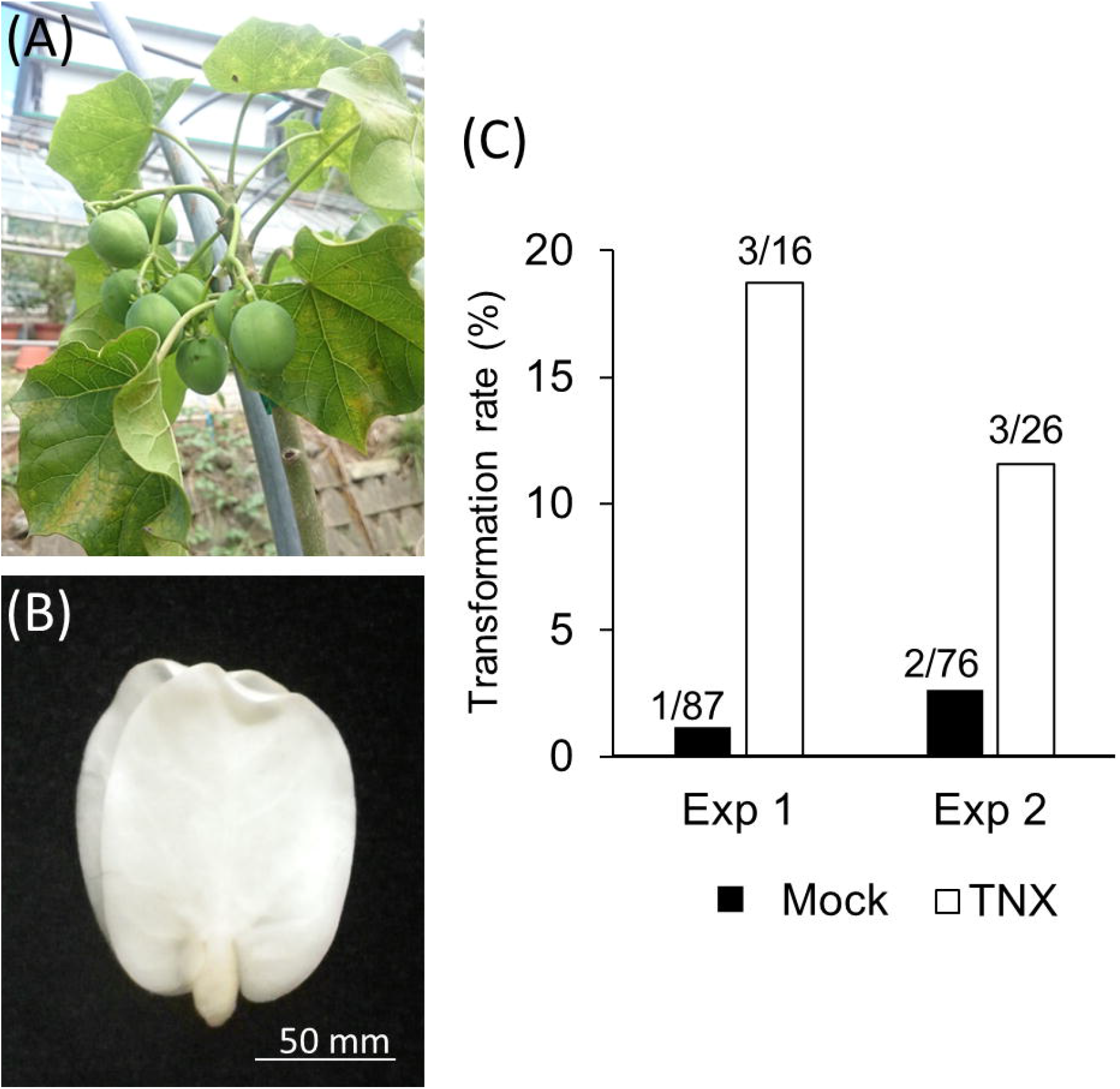
Tenoxicam (TNX) treatment enhances the efficiency of *Agrobacterium*-mediated stable transformation in Jatropha. (A and B) Images of mature Jatropha plants (A) and the cotyledon used for transformation (B). Bar, 50 mm. (C) Transformation rate determined by PCR genotyping in the mock- or TNX (100 μM)-treated plants. The data obtained from two independent experiments are shown. Numbers above the bars indicate number of transformants with respect to the number of treated soil-acclimated plants.

**Figure 4.**
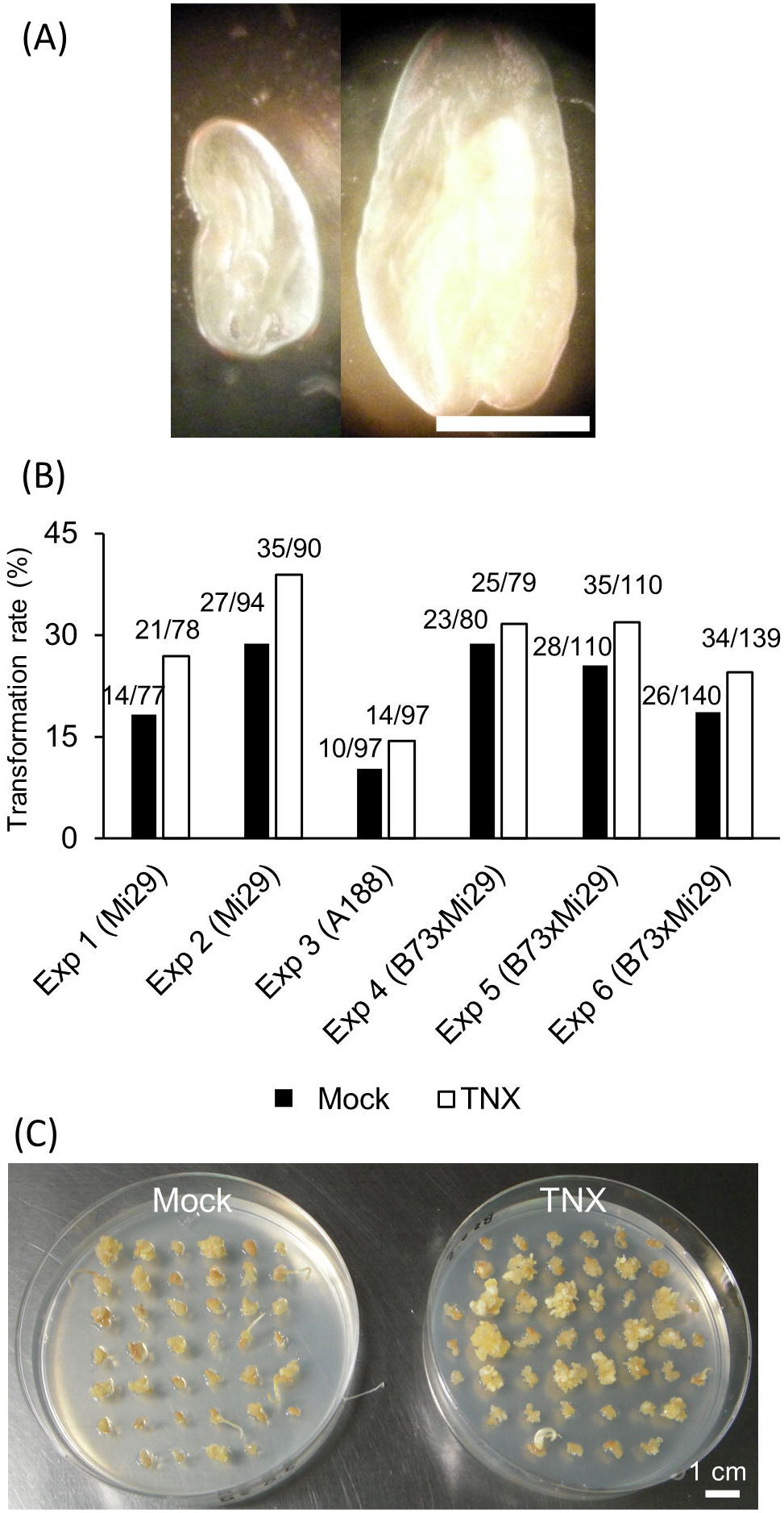
Tenoxicam (TNX) treatment enhances the efficiency of stable *Agrobacterium*-mediated transformation in maize. (A) Image of the immature embryo of maize that was used for transformation. Bar, 1 mm. (B) Transformation rate determined based on survival rate on selection medium for mock- and TNX (100 μM)-treated plants. The data obtained from six independent experiments using different strains are shown. (C) Image of calli on selection media after mock (left) and 100 μM TNX (right) treatments. Bar = 1 cm.

Maize is an important food in many countries and has a range of biotechnological-related traits of commercial interest. Accordingly, maize is considered one of the most important target crops for the application of biotechnology (Que et al., 2014). In the present study, we performed transformation of maize using immature embryos (Fig. 4A) by adding TNX to the co-cultivation suspension or medium in six independent experiments using different maize strains. Using selection medium, we assessed the survival rate of callus derived from the TNX-treated and non-treated immature embryos, and in all independent experiments observed survival rates in the TNX treatments that were comparable to or higher than those obtained in the control treatments, particularly with respect to the Mi29 strain, which showed high sensitivity to TNX in terms of transformation efficiency (Fig. 4B). Surprisingly, we found that the TNX-treated calli tended to grow larger (Fig. 4C), thereby indicating that a larger number of cells within the immature embryos had been transformed in response to the TNX treatment to form a larger callus mass.

### Meristematic tissues are insensitive to tenoxicam

We also evaluated the effect of TNX treatment on the efficiency of transformation for other plant species, namely rice, soybean, *Brassica napus*, *Brassica rapa*, and water starwort. Unfortunately, we failed to detect a significant effect of TNX treatment on the efficiency of transformation for these plants (Fig. S1, Table S3-S6). However, with the exception of water starwort, we used meristematic tissues of these plant, such as callus, callus-induced tissues, or cotyledonary nodes, rather than mature vegetative tissues, thereby indicating that the effect of TNX may be tissue dependent. In this context, it is well established that plant growth and immunity show antagonistic interactions; for example, young tissues must suppress the immune response to maximise growth in the absence of perceived pathogens, whereas mature organs are more adapted for defensive roles (Wang and Wang, 2014). This is supported by the different defence-related responses observed during different phases of the cell cycle (Kadota et al., 2004). In this regard, SA has been established to be a key regulator promoting immunity but suppressing growth, whereas the atypical E2F protein DEL1 promotes cell proliferation and suppresses expression of the SA transporter gene *enhanced disease susceptibility 5* (*EDS5*) and the SA biosynthetic gene *ISOCHORISMATE SYNTHASE1* (*ICS1*) to suppress SA accumulation and defence responses in growing tissues (Chandran et al., 2014). Given that meristematic tissues already show less pronounced immune responses and that TNX can inhibit immune signalling pathways, including the SA pathway, we speculate that these meristematic tissues could be insensitive to TNX. This would be consistent with our observations that *EDS5* and *ICS1* are highly expressed in the mature tissues of *A. thaliana*, such as leaves, whereas expression is lower in meristematic tissues (Figure S2), which would explain why TNX, although efficient with respect to leaf agroinfiltration, appears to be ineffective for the transformation of flowers and roots (Fig. 2 and Tables S1 and S2).

### Conclusions

In this study, we demonstrated that oxicam-type NSAIDs, including tenoxicam, can enhance the efficiency of *Agrobacterium*-mediated transformation in *A. thaliana*, as well as in certain crops of agricultural and economic importance, namely, Jatropha and maize. Given that treatment with oxicam compounds is comparatively straightforward (simply adding chemicals to the *Agrobacterium* co-cultivation medium), cost effective (estimate one US dollar for one hundred transformants), and has broad utility (it can readily be applied using existing methods), we anticipate that this discovery will contribute to research applications such as transient transformation assays, as well as potentially providing a solution for worldwide food security problems. However, we found that TNX had no appreciable effect with respect to the efficiency of meristematic tissue transformation, which we suspect might be attributable to the insensitivity of such tissues to TNX, given that the immune responses of meristematic tissues are typically suppressed. Therefore, we propose that future research in this area should focus on the monitoring and modification of plant immune responses, which we believe would provide a basis for overcoming some of the current limitations associated with plant transformation.

## Materials and Methods

### Arabidopsis thaliana

For the purposes of agroinfiltration, we used 5- to 6-week-old Col-0 and *dde2/ein2/pad4/sid2* quadruple mutant *Arabidopsis* plants grown in soil under short-day conditions (8/16 h light/dark). *Agrobacterium tumefaciens* (strain Agl1) carrying the GUS reporter gene in a pGreen0029 vector used for infiltration was prepared and resuspended at an OD_600_ of 0.2, as previously described (Choi et al., 2013). For co-expression of organellar markers, *A. tumefaciens* (strain GV3101) transformants harbouring *RPS::H2B-GFP*, *35S::Venus-SYP61*, and *35S::PIP2A-mCherry* were mixed after resuspension. Oxicams [tenoxicam (T0909; Sigma-Aldrich), meloxicam (M3935), piroxicam (P5654), ampiroxicam (SML1475), lornoxicam (SML0338), and sudoxicam (S688950; Wako Chemicals) dissolved in DMSO] were added to a final concentration of 100 μM in the resuspension medium immediately prior to infiltration. Within 48 h, GUS activity was measured using MUG assays and subcellular localisation was observed under a confocal microscope.

For histochemical staining to visualise GUS activity, we used 5-bromo-4-chloro-3-indolyl-β-D-glucuronide (X-gluc) as a substrate. Infiltrated leaves were collected and fixed in 90% acetone on ice for 15 min, after which the samples were placed in GUS staining solution [1 mg/ml X-gluc, 100 mM NaPO_4_(pH 7), 0.5 mM K_3_Fe(CN)_6_, 0.5 mM C_6_N_6_FeK_4_, 10 mM EDTA, and 0.1% triton X-100] under vacuum at room temperature for 15 min and incubated overnight at 37°C. Subsequent to GUS detection, leaves were rinsed with 70% ethanol and cleared in chloral hydrate:water (2.5:1).

For quantitative MUG assays, leaf discs were homogenised in extraction buffer (50 mM NaHPO_4_ pH 7.0, 10 mM EDTA, 0.1% sarcosyl, 0.1% Triton X-100, and 10 mM β-mercaptoethanol) and centrifuged at 4°C. Ten microlitres of the supernatant was added to 90 μL of assay buffer [2 mM 4-methylumbelliferyl-β-D-glucuronic acid (MUG) in extraction buffer without β-mercaptoethanol] in wells of a 96-well microplate and incubated at 37°C in the dark for 30 min. Aliquots (10 μL) of the reaction mixtures were mixed with 90 μL of Na_2_CO_3_ and the fluorescence of 4-methylumbelliferone (4-MU) was measured at excitation at 365 nm and emission at 455 nm using a Mithras LB940 multiplate reader (Berthold Technologies). GUS activity was calculated as pmol 4-MU formed per mg total protein per min.

Subcellular localisation was observed using a Zeiss LSM 700 inverted confocal microscope. GFP and Venus were excited at 488 nm, and emission was recorded between 501 and 545 nm, whereas mRFP was excited at 561 nm and the emission was recorded between 570 and 615 nm.

For root transformation, Col-0 plants were grown for more than 14 days on a plate containing Murashige and Skoog (MS) salt mixture (Wako Pure Chemical Industries, Ltd, Osaka, Japan) under long-day conditions (16/8 h light/dark). Roots were cut into 5-mm segments, and bundles of roots were inoculated with *A. tumefaciens* (GV3010) carrying *35S: GFP* on an MS plate. Prior to application, DMSO or 100 μM TNX was added to the *Agrobacterium* suspension. After 2 days of co-cultivation, transiently expressed GFP signals were observed from root bundle tips, and the number of tips expressing GFP was counted. The root bundles were then transferred to callus-inducing medium [CIM: 1× MS, 0.05 M MES, vitamins (0.5 mg/L nicotinic acid, 0.5 mg/L pyridoxine, 0.5 mg/L thiamine-HCl), 100 mg/L myo-inositol, 20 g/L glucose, 5 mg/L indole-3-acetic acid (IAA), 0.5 mg/L 2,4-dichlorophenoxy acetic acid (2,4-D), 0.3 mg/L kinetin, pH 5.7, and 7.5 g/L Bacto agar]. After 4 weeks, callus had developed from all root bundles, and the number of GFP signal spots on callus. detected using a Leica M165 C stereomicroscope, was counted.

For floral-drop transformation, *A. tumefaciens* (strain Agl1) harbouring *35S::GUS* was grown overnight at 28°C in 30 mL of Luria–Bertani (LB) medium containing rifampicin and kanamycin. Cells were grown to the stationary phase (an OD_600_ of approximately 2.0) and harvested by centrifugation at 4500 × *g* for 10 min. The pellet was resuspended in medium (0.5× MS, 5% sucrose, and 0.05% Silwett L-77) and DMSO or 100 μM TNX was added to the *Agrobacterium* suspension. We performed drop-wise inoculation of individual flower buds using a micropipette. Seeds collected from inoculated plants were sown on medium supplemented with 50 mg/L kanamycin and 100 mg/L cefotaxime sodium and incubated for approximately 10 days to assess kanamycin resistance.

### Jatropha

In this study, we transformed Jatropha (*Jatropha curcas* L.) using *A. tumefaciens* (strain EHA101) harbouring selection vectors (PalSelect A-3 vector, Kumiai Chemical Industry Co., Ltd.). The transformation procedure used has been described previously by Enoki et al. (2017). Briefly, 3–5-mm square segments of cotyledons (Fig. 2B) were co-cultivated with *Agrobacterium* suspension supplemented with DMSO or 100 μM TNX at 26°C for 4 days in the dark. Calli and shoots that had developed after culturing on CIM followed by shoot induction medium, were transferred to selection medium. Elongated shoots were subsequently transferred to root induction medium and when roots had become fully developed, the transformed plants were transferred to soil and acclimatised in a growth room until genotyping by PCR using primers specific for the transgene.

### Maize

For the transformation of maize (*Zea mays* L.), we used the inbred line Mi29 and *A. tumefaciens* (strain LBA4404) harbouring a pVGW9 helper plasmid carrying the bar gene driven by a ubiquitin promoter. The method used for transformation has been described previously by Ishida et al. (2007). Immature embryos obtained at approximately 10 to 18 days (depending on the season) after pollination, were co-cultivated on LSAS medium containing DMSO or 100 μM TNX at 25°C for 7 days. The embryos were then transferred to LSD1.5A medium containing 3 mg/mL bialaphos, 250 mg/L carbenicillin, and 100 mg/L cefotaxime and incubated for 10–20 days.

### Rice

For rice (*Oryza sativa* L.) transformation, we used the cultivar Nipponbare and *A. tumefaciens* (strain EHA105, Hood et al., 1993) harbouring a monitoring vector (Saika et al., 2012). The transformation procedure followed that described by Toki (1997). Briefly, 1-week-old primary calli were co-cultivated with an *Agrobacterium* suspension supplemented with DMSO or 100 μM TNX at 23°C for 3 days. *Agrobacterium*-infected calli were subsequently cultured on selection medium containing 10 mg/L blasticidin S (Nacalai Tesque) and 25 mg/L meropenem (Wako Pure Chemical Industries) at 31–33°C. Transformed calli were treated with Beetle Luciferin, potassium salt (Promega) and left to stand for 5 min. Thereafter, Luc luminescence images were obtained and analysed using a Lumazone imaging system (Roper Scientific).

### Soybeans

For the purpose of soybean [*Glycine max* (L.) Merr.] transformation, we used the Japanese soybean variety Kariyutaka, which was obtained from the Hokkaido Prefectural Agricultural Experiment Station, Tokachi, and is identical to the resource JP 86520 available from the Genebank Project, National Agriculture and Food Research Organization. *Agrobacterium*-mediated transformation was performed using *A. tumefaciens* (strain EHA105) as previously described by Yamada et al. (2014). Either DMSO or 20 μM TNX was added to the co-cultivation medium. Integration of the transgene was assessed by applying Basta (Bayer Crop Science) to the leaves of T_1_ seedlings.

#### Brassica napus

*Brassica napus* cv. Westar was transformed by inoculating hypocotyl sections with *A. tumefaciens* (strain GV3101) harbouring a binary vector containing the neomycin phosphotransferase II gene and regenerated as described by Kohno-Murase et al. (1994). DMSO or 100 μM TNX was added to the co-cultivation medium. The numbers of transformed plants grown on selection medium were counted.

#### Brassica rapa

The seeds of *Brassica rapa* L. subsp. *nipposinica* (‘Kyo-nishiki’ cultivar of Mibuna) used for transformation were purchased from Takii & Co., Ltd., Kyoto, Japan. These seeds were sterilised in 70% ethanol for 2 min, soaked and swirled in 1% sodium hypochlorite for 15 min, and then washed five times with sterile distilled water to remove the sodium hypochlorite. The surface-sterilised seeds were poured into a magenta box containing germination medium [1× MS medium (Wako, Japan) containing 3% sucrose, 0.1% PPM (Plant Cell Technology, USA), and 0.4% Gelrite (Wako, Japan)], and incubated at 20°C for 7 days for germination. Hypocotyl explants from the germinated plants were subsequently pre-cultured for 3 days on CIM [B5 salt and vitamins (Wako), 1.5% sucrose, 0.2% PPM, and 1 mg/L 2-4D, pH 5.8)]. *A. tumefaciens* (strain C58C1) carrying the binary vector pBCR101 was cultured overnight at 28°C in liquid LB medium containing 50 mg/L kanamycin. The following day, the *Agrobacterium* cells were harvested by centrifugation at 8,000 rpm for 20 min and then re-suspended to an OD_600_ of 0.1 with infection medium (Liquid MS, pH 5.8, and 200 μM acetosyringone) containing DMSO or 100 μM TNX. The pre-cultured hypocotyl explants were soaked in the bacterial suspension for 10 min and then transferred to sterile paper to remove the *Agrobacterium*. The infected hypocotyl explants were then placed on CIM and incubated at 20°C for 3 days in the dark. Following co-cultivation, the explants were washed with sterile distilled water and cultured on CIM containing 200 mg/L carbenicillin for 7 days, and subsequently transferred to fresh CIM at 2-week intervals. After one and half months, calli had formed from the hypocotyl explants. As we were unable to obtain sufficient redifferentiation using the Mibuna cultivar, transformation efficiency was assessed by GUS staining of callus that developed from the hypocotyl explants.

### Water starwort

Seeds of water starwort (*Callitriche palustris* L.), collected from a laboratory culture strain (originally collected from Nagano, Japan) were sterilised with 4% sodium hypochlorite solution (Wako, Japan) for 20 min and then incubated in sterile distilled water for 3–7 days to induce germination. The germinated seedlings were thereafter transplanted onto germination medium [0.5× MS medium (Wako, Japan) containing 2% sucrose and 0.3% Gellan Gum (Wako, Japan)], and cultured in a growth chamber (NK system, Osaka, Japan) at 23°C under constant light at an illumination intensity of 60 μmol m^-2^ s^-1^. Immediately prior to transfection, hypocotyls were dissected from 2-week-old seedlings as explants. *A. tumefaciens* (strain C58C1Rifr) harbouring a pH35G binary vector, in which a cDNA sequence for GFP was linked to the *35S CaMV* promoter, were cultured in LB medium supplemented with 50 mg/L ampicillin and 100 mg/L spectinomycin at 28°C for 24 h. For transfection, the *Agrobacterium* was resuspended in 0.5× MS supplemented with 2% sucrose. The explants were immersed in the *Agrobacterium* suspension containing DMSO or 50 μM or 100 μM TNX in addition to 100 μM acetosyringone, and then incubated for 10 min. The explants were thereafter transferred onto a solid medium consisting of 0.5× MS, 2% sucrose, 2 mg/L *trans*-zeatin (Tokyo Chemical Industry Co., LTD, Toyo, Japan), and 100 μM acetosyringone. After co-cultivation for 3 days at 23°C in the dark, the *Agrobacterium* was removed by incubating the explants on 0.5× MS medium containing 250 mg/L cefotaxime sodium salt (Sanofi K.K., Tokyo, Japan), 2% sucrose, and 2 mg/L *trans*-zeatin for 2 days under illuminated growth conditions. The numbers of explants containing GFP-positive cells were counted under an MZ16 fluorescence stereomicroscope (Leica Microsystems, Wetzlar, Germany).

## List of abbreviations

## Author contributions

S-WC, KS and YI conceived and designed the experiments. KK, RM, HE, WC, TT, HS, ME, TY, AH, NK, SK, YK, and HK performed the experiments. YI wrote the first draft of the manuscript. S-WC, EI, KS, and YI reviewed and edited the manuscript. All authors contributed to the article and approved the submitted version.

## Acknowledgements

We thank Dr. Shuta Asai (Riken Center for Sustainable Resource Science) for providing seeds of the *dde2/ein2/pad4/sid2* quadruple mutant, Dr. Hirokazu Tsukaya (The University of Tokyo) for providing helpful advice regarding the experiments using water starwort, and Dr. Goro Horiguchi (Rikkyo University) for providing the binary vector pH35G.

