## Supplemental data for "Oxicam-type NSAIDs enhance *Agrobacterium*-mediated transformation in plants"

### *Supplementary Material*

#### 1.1 Supplementary Figures

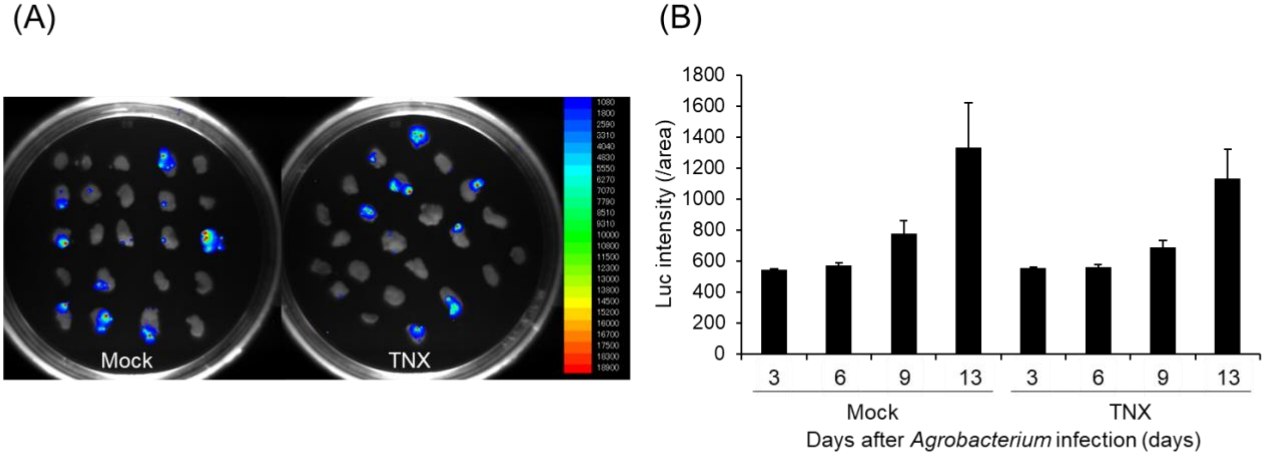


**Supplementary Figure 1. Effect of tenoxicam (TNX) on the *Agrobacterium*-mediated transformation of rice** (A) Luminescence emitted from callus transformed with the monitoring vector at 13 days after *Agrobacterium* infection in mock- or TNX (100 µM)-treated tissue. Relative luminescence intensity is shown by the false colour scale. (B) Quantification of luc intensity at 3, 6, 9, and 13 days after *Agrobacterium* infection to compare the mock and 100 µM TNX treatments. Values are the average ± SE (n = 23).

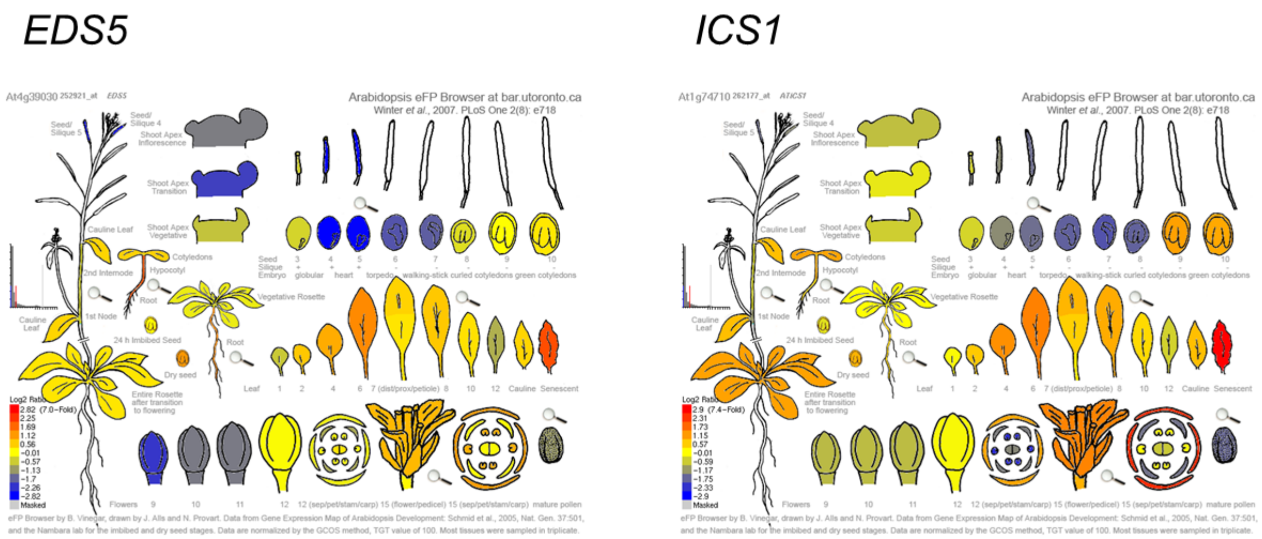


**Supplementary Figure 2. *EDS5* and *ICS1* gene expression in different tissues of *Arabidopsis thaliana.*** The relative gene expression patterns of *EDS5* (AT4G39030) and *ICS1* (AT1G74710) are shown. The images were obtained from the eFP browser (http://bar.utoronto.ca/efp/cgi-bin/efpWeb.cgi) showing a comprehensive gene expression map for different tissues at different stages of development.

#### 1.2 Supplementary Tables

**Supplementary Table 1. Effect of tenoxicam (TNX) on the *Agrobacterium*-mediated root transformation in *Arabidopsis thaliana***

| Experiment | TNX conc. (μM) | Transient expression | | | Number of GFP spots on callus |
|---|---|---|---|---|---|
| | | Total number of root bundle tips | Number of root bundle tips showing GFP signals | Transformation rate (%) | |
| 1 | 0 | 268 | 34 | 12.7 | 36 |
| | 50 | 316 | 84 | 26.6 | 61 |
| | 100 | 299 | 49 | 16.4 | 43 |
| 2 | 0 | 375 | 115 | 30.7 | 96 |
| | 50 | 177 | 84 | 47.5 | 71 |
| | 100 | 371 | 117 | 31.5 | 112 |
| 3 | 0 | 351 | 54 | 15.4 | 53 |
| | 50 | 348 | 79 | 22.7 | 46 |
| | 100 | 337 | 67 | 19.9 | 68 |
| 4 | 0 | 353 | 93 | 26.3 | 70 |
| | 50 | 259 | 102 | 39.4 | 76 |
| | 100 | 343 | 100 | 29.2 | 103 |
| 5 | 0 | 217 | 36 | 16.6 | 42 |

| | | | | |
|---|---|---|---|---|
| 50 | 207 | 27 | 13.0 | 30 |
| 100 | 294 | 55 | 18.7 | 45 |

**Supplementary Table 2. Effect of tenoxicam (TNX) on the *Agrobacterium*-mediated floral-drop transformation in *Arabidopsis thaliana***

| Experiment | TNX conc. (μM) | Number of $T_0$ seeds | Number of seeds resistant to antibiotics | Transformation rate (%) |
|---|---|---|---|---|
| 1 | 0 | 1,000 | 5 | 0.5 |
| | 100 | 1,000 | 8 | 0.8 |
| 2 | 0 | 1,000 | 3 | 0.3 |
| | 100 | 1,000 | 5 | 0.5 |
| 3 | 0 | 1,000 | 7 | 0.7 |
| | 100 | 1,000 | 3 | 0.3 |

**Supplementary Table 3. Effect of tenoxicam (TNX) on the *Agrobacterium*-mediated transformation of soybean**

| Experiment | TNX conc. (μM) | Total number of explants | Explants generating $T_0$ plants | | Explants generating stable $T_0$ plants | |
|---|---|---|---|---|---|---|
| | | | Total number | Transformation rate (%) | Number of Transformed $T_0$ | Transformation rate (%) |
| 1 | 0 | 86 | 3 | 3.5 | 1 | 1.2 |
| | 20 | 88 | 4 | 4.5 | 1 | 1.1 |
| 2 | 0 | 90 | 4 | 4.4 | 1 | 1.1 |
| | 20 | 101 | 8 | 7.9 | 2 | 2.0 |
| 3 | 0 | 110 | 5 | 4.5 | 1 | 0.9 |
| | 20 | 110 | 3 | 2.7 | 1 | 0.9 |
| 4 | 0 | 60 | 10 | 16.7 | 1 | 1.7 |
| | 20 | 120 | 12 | 10.0 | 3 | 2.5 |

**Supplementary Table 4. Effect of tenoxicam (TNX) on the *Agrobacterium*-mediated transformation of *Brassica napus***

| Experiment | TNX conc. (μM) | Total number of explants | Explants generating $T_0$ plants | |
|---|---|---|---|---|
| | | | Total number | Transformation rate (%) |
| 1 | 0 | 338 | 8 | 2.4 |
| | 100 | 337 | 7 | 2.1 |
| 2 | 0 | 198 | 1 | 0.5 |
| | 100 | 198 | 0 | 0.0 |
| 3 | 0 | 179 | 1 | 0.6 |
| | 100 | 180 | 1 | 0.6 |
| 4 | 0 | 208 | 1 | 0.5 |
| | 100 | 208 | 1 | 0.5 |

**Supplementary Table 5. Effect of tenoxicam (TNX) on the *Agrobacterium*-mediated transformation of *Brassica rapa***

| Experiment | TNX conc. (μM) | Average number of GUS spots on explants (±SE) |
|---|---|---|
| 1 | 0 | 2.3 (±0.4) n = 18 |
| | 100 | 1.1 (±0.7), n = 7 |
| 2 | 0 | 1.7 (±0.3), n = 29 |
| | 100 | 1 (±0.6), n = 7 |

**Supplementary Table 6. Effect of tenoxicam (TNX) on the *Agrobacterium*-mediated transformation of *Callitriche palustris***

| Experiment | TNX conc. (μM) | Total number of explants | Explants with transformed cells | |
|---|---|---|---|---|
| | | | Total number | Transformation rate |
| 1 | 0 | 39 | 31 | 0.8 |
| | 50 | 30 | 20 | 0.7 |
| | 100 | 30 | 21 | 0.7 |
| 2 | 0 | 39 | 9 | 0.2 |
| | 50 | 38 | 8 | 0.2 |
| | 100 | 42 | 17 | 0.4 |
| 3 | 0 | 35 | 24 | 0.4 |
| | 50 | 44 | 24 | 0.5 |
| | 100 | 45 | 25 | 0.6 |
| 4 | 0 | 37 | 27 | 0.7 |
| | 50 | 40 | 28 | 0.7 |
| | 100 | 37 | 27 | 0.7 |
